# What doesn’t kill you makes you stronger: detoxification ability as an honest sexually selected signal

**DOI:** 10.1101/2020.05.09.086280

**Authors:** Isaac González-Santoyo, Daniel González-Tokman, Miguel Tapia-Rodríguez, Alex Córdoba-Aguilar

## Abstract

Sexual selection maintains colourful signals that increase sexual attractiveness and dominance. Some sexually selected, colourful signals are pigments synthesized from ingested amino acids. The underlying metabolic pathways for these pigments often release toxic byproducts that can reduce individual survival. However, rather than discarding these otherwise harmful byproducts, animals may use them by integrating them into sexually selected traits. We tested this idea using males of the damselfly *Hetaerina americana*, which bear a red-pigmented wing spot that is sexually selected through male-male competition for mating territories. First, by using chromatography and confocal microscopy, we determined that the red wing spots are generated by ommochrome pigments derived from tryptophan metabolism. Second, we injected a group of males with the toxic precursor of these ommochromes, 3-hydroxy-kynurenine (3-Hk), confirming the toxicity of this compound in adult males. Finally, by using spectrophotometry and confocal microscopy, we showed that adult males injected with a LC_50_ of 3-Hk had more ommochromes in their wing spots than controls but similar survival, suggesting that the deposition of ommochrome pigment in the wing detoxifies the tryptophan metabolism process. Thus, we report for the first time that sexually selected pigmented signals involve the biochemical treatment of excreted compounds that could otherwise have lethal effects, a hypothesis we call “detoxifying ability signalling”. Our results provide new insights about the origin and maintenance of sexual signals, elucidating a mechanism for the evolution of honest indicators of quality that could have arisen due to natural selection.

## Introduction

Some animal colours result from the sequestration as pigments in the integument of harmful metabolic products that cannot be easily eliminated from the body during digestion ^1^. This mechanism is called storage excretion. What makes this mechanism particularly interesting is that rather than being harmful, the excreted pigments can generate external colouration that has adaptive functions such as camouflage or signalling information to conspecifics and heterospecifics. For example, excessive uric acid, the main nitrogen metabolic waste of terrestrial insects, sometimes cannot be completely eliminated during digestion but is excreted as a white or yellow pigment that is deposited in the larval cuticle of several species of pierid butterflies ^2^. This deposition creates the impression of bird droppings, which looks distasteful for predators, decreasing predation and thus increasing larval survival ^2^. Similar pigmentation patterns in adult butterflies, resulting from storage excretion, have been proposed to be involved in sexual selection processes, such as colourful traits used for attracting mates or dissuading conspecifics during male-male competition.

A classical tenet in sexual selection theory is that the expression of pigmented traits has evolved because these colours grant individuals an advantage during mate choice or intrasexual competition for mates ^3^. For instance, it has been observed that bright male colouration is maintained by sexual selection because it can only be produced by high quality (e.g. well-fed) individuals that can afford the production of pigments without compromising other traits that require the same limiting resource ^4^. Interestingly, genes responsible for colouration interact pleiotropically with other genes that can be involved in important physiological functions such as thermoregulation, photoprotection, desiccation resistance, immunoregulation, antioxidation and excretion of toxic compounds that may result from metabolism and immune response ^5–7^. Therefore, pigmentation that reflects efficient metabolism may be subject to strong sexual selection if they honestly reflect individual genetic and physiological condition ^6,8^.

Storage excretion as a mechanism of sexual pigmentation has not been discussed and examined in detail despite providing a clear physiological and evolutionary explanation of the origin and maintenance of many sexual signals. Storage excretion can be particularly important when the synthesis of such pigments is involved in physiological pathways that generate waste products that are difficult to eliminate or inactivate. Moreover, the assumption that the trait provides honest information about the quality of the bearer is upheld, since the use of toxic byproducts to generate visible colouration could be doubly informative: pigments signal the bearer’s ability to deal with toxic metabolites that could otherwise impair their survival, as well as the genetic and/or physiological condition since for example, because only individuals with better diets or protein metabolism will produce a large enough amount of the byproducts to require storage excretion of the pigments (as suggested by ^6^).

Red pigments are common in several invertebrates, and most of their underlying components are ommochromes ^9^. In insects, ommochromes give colour to structures such as the eyes and the wings of flies, butterflies and damselflies ^10–12^. These red pigments result from the metabolism of tryptophan, an essential amino acid that is consumed from food ^12,13^. Ommochrome pigments are synthetized from the tryptophan metabolite 3-hydroxykynurenine (3-Hk), which together with tryptophan, are considered neurotoxic compounds that cause paralysis, altered mating behaviour, aging and high mortality in adult insects ^14–17^. In fact, wild type flies fed with inhibitors of tryptophan-kynurenine metabolism show increased survival compared to control flies ^17^. 3-Hk is present at high concentrations during metamorphosis when protein breakdown releases high quantities of this tryptophan product ^12^. Given the noxious effects of 3-Hk, an important way in which insects can deal with excessive neurotoxic byproducts is via the synthesis of ommochromes, which can be eliminated in the meconium during pupation in holometabolous insects, in the excreta or by storage excretion in the form of cuticular pigmentations, a hypothesis proposed by Linzen in 1974 and called “the tryptophan-detoxification hypothesis” ^5,9,18^. In fact, both 3-Hk and ommochrome pigments are found at high concentrations in the meconium of lepidopterans, the waste product from the pupal stage ^19^.

Ommochromes can function as sexually selected signals and may be considered honest indicators of individual quality, since only well-fed individuals will accumulated high enough amounts of these metabolites to develop conspicuous signals ^20^.

Here, we propose that ommochromes as component of male sexual traits can be used for detoxification, providing a new mechanism of sexual signal production and fulfilling the honesty principle. We tested these ideas using males of the rubyspot damselfly *Hetaerina americana*. This species is a classic model in sexual selection studies: males bear a red wing spot (RWS; Fig. 1a) whose size is an honest indicator of individual condition, and which provides an advantage during male-male competition ^21,22^. We (1) determined the presence of ommochromes in the wing spot, (2) tested the toxic effect of 3-hydroxykynurerine (3-Hk), the most toxic tryptophan metabolite and precursor of ommochromes, and (3) measured deposition of ommochromes in wing spots of young adult males injected with 3-Hk.

**Figure 1.**
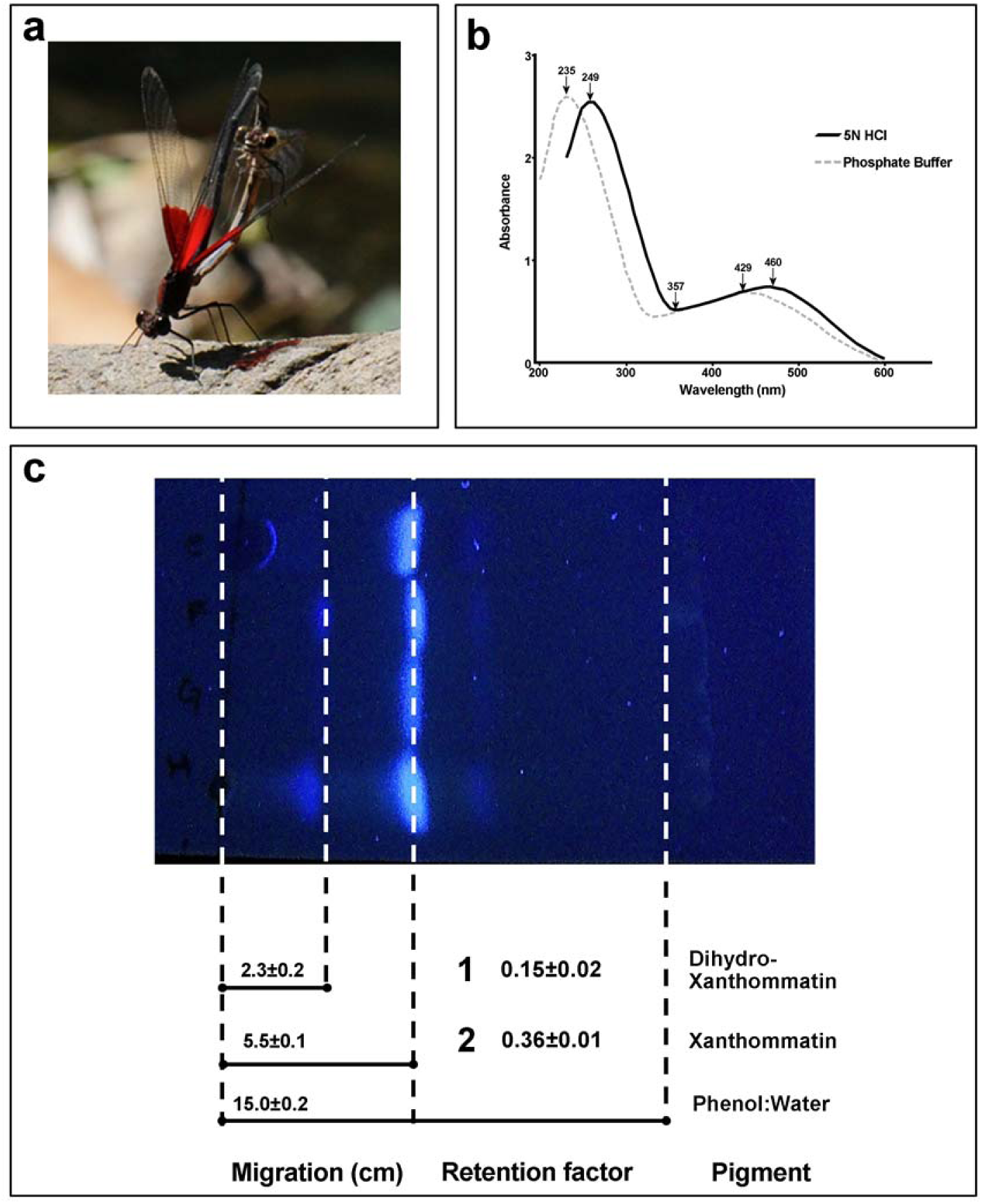
(a) *Hetaerina americana* male showing his red wing spot (RWS) during mating. (b) Spectral absorbance analysis showing the presence of ommochrome pigments in the RWS. The red powder dissolved in 5N HCl (solid black line) or phosphate buffer (dashed grey line) show a typical spectral behaviour of the pigment xanthommatin. (c) UV spectrum photography from the thin layer chromatography (TLC) confirmed the presence of xanthommatin. The column migration and dashed lines show the distance travelled (cm) by the two bands observed in the TLC and by the solvent used (Phenol:Water). The retention factor (Rf) for these bands (1 and 2) were consistent with the authentic standard of xanthommatin (number 2) and its reduced form dihydro-xanthommatin (number 1). The Rf of the authentic standard of xanthommatin was reported by Nijouth (1997) using the same TLC method.

## Results

All biochemical test confirmed the presence of ommochromes as main pigments in the RWS (Fig. 1a). Besides Redox Behaviour, we observed that the absorption spectra from the red powder diluted in 5N HCl displayed three absorbance wavelength peaks; 249, 357 and 460 nm with relative absorbances of 2.49, 0.52 and 0.74 respectively. The red pigment dissolved in phosphate buffer (pH 7.0) displayed peaks at two different wavelengths: 235 and 429 nm with relative absorbances of 2.68 and 0.68 respectively (Fig. 1b).

Moreover, our TLC analysis indicates that xanthommatin is an ommochome present in *H. americana* male RWS. When the silica gel plate was exposed to the UV spectrum, we observed a slightly defined band with a *Rf* = 0.15+0.013 and a well-defined band with *Rf* = 0.36+0.012 (Fig. 1b), which coincides with the standard synthetic ommochrome xanthommatin (*Rf=*0.36) when the same silica gel/phenol TLC system is implemented ^23^. The other slightly defined band observed showed a similar *Rf* value to the same xanthommatin, but in its reduced form, dihydro-xanthommatin (Rf=0.13 ^23^). We also found a slight third band (Fig. 1b), which may correspond to 3-Hk (Rf=0.52 ^23^), although it was very faint.

In addition, patterns of autofluorescence in the RWS were observed in the spectral confocal microscope. Figure 2 shows MIP images obtained from XYZλ scans of both red (Fig. 2a) and transparent (Fig. 2b) regions of a representative control wing (Fig. 2c) excited with 561 nm wavelength light. We excited separately with 405, 488, 561 and 647 nm wavelengths and found two main types of fluorescence: the most conspicuous corresponds to ommochrome pigments, which are distributed along the wing veins (Fig. 2a and d) and spread over the wing tissue in the red pigmented area (Fig. 2d and 2e), but not in the transparent region (Fig. 2b). This signal was strongest when excited at 561 nm, but it was also visible when excited at 405 or 488 but not at 647 nm wavelengths (see supplementary figures 1a, 1b and 1c). The other main type of fluorescence observed was autofluorescence of both the red and the transparent wing regions, which can be attributed to the reflection of light by the wax that covers the wings. This type of fluorescence occurred at all wavelengths (see supplementary figures 1a, 1b and 1c) and shows a characteristic periodicity on its distribution pattern.

**Figure 2.**
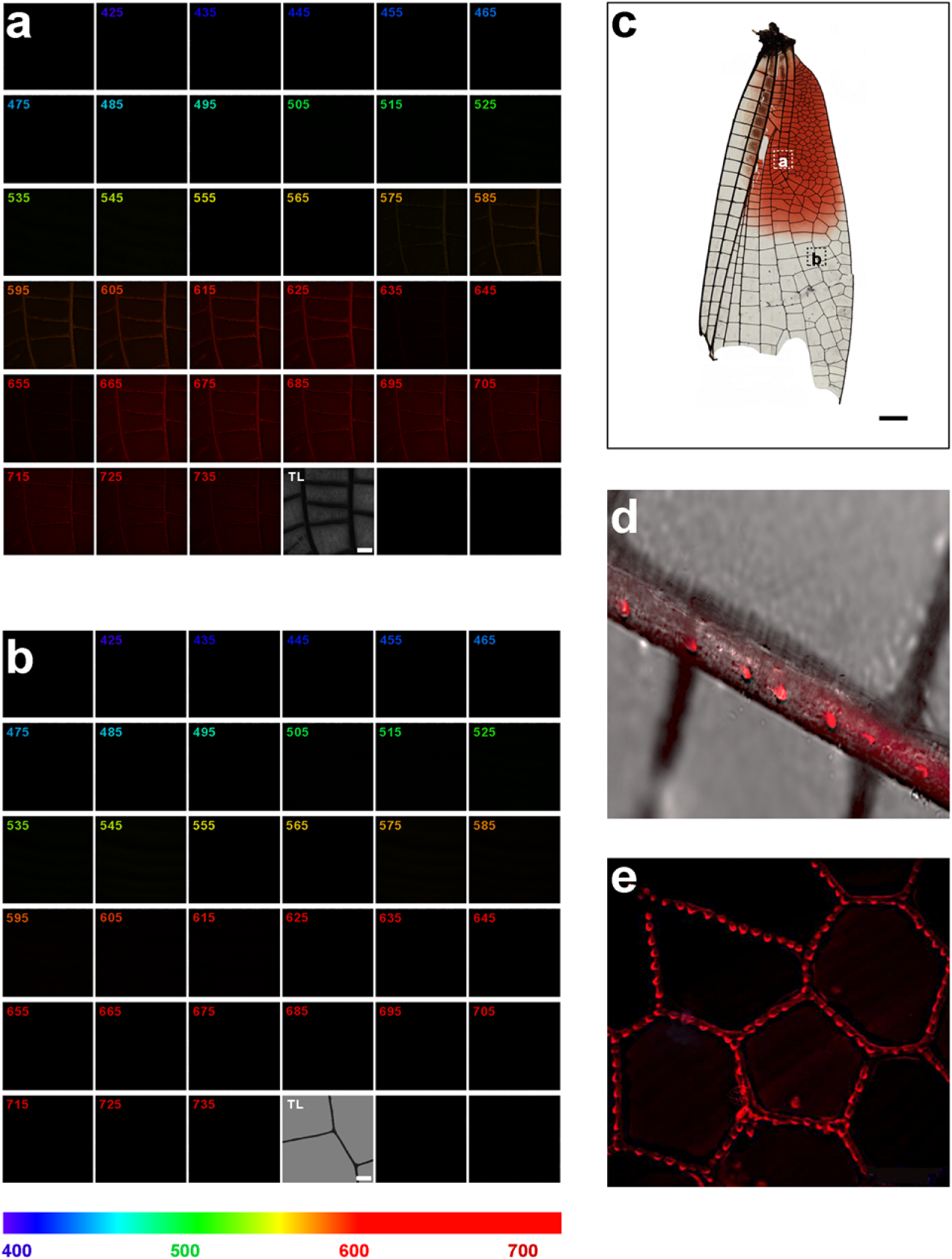
Fluorescence XYZλ scans (425-735 nm) of *H. americana* wings excited at 561 nm. Maximum intensity projection images of (a) a sub-region of the red-pigmented area, and (b) a sub-region of the transparent area, depicted in (c). Each frame in (a) and (b) represent a specific wavelength image, as indicated. Fluorescence intensity was highest between 615-725 nm in the red pigmented area but not in the transparent region. Higher magnifications show the presence of red fluorescence spots distributed along the wing veins ç(d) and diffusely spread over the wing tissue (e). Scale bars in a and b indicate 100μm and in c, 1mm.

3-Hk had a range of toxic effects on *H. americana* males, from partial immobility at the lowest concentrations tested (1 and 100 μg mL^−1^) to death for the highest concentrations (1000 and 10000 μg mL^−1^). This toxicity led us to determine the LC_50_ for sexually mature adult males. We found that the group with the lowest 3-Hk concentration (1 μg mL^−1^) showed a mortality rate of 13.13%, while subsequent doses (100, 1000 and 10000 μg mL^−1^) increased mortality to 40%, 46% and 53% respectively. Consequently, LC_50_ of 3-Hk for males of this species was estimated at 368.69 μg mL^−1^ (C.I 95%: 78.5-1729.9μg mL^−1^).

The mitigation of 3-Hk toxic effects by deposition of ommochromes in RWS was observed in males treated with the LC_50_ of 3-Hk, since they obtained similar survival to sham males (z=1.38, P=0.167; Figure 3); Control males survived less than sham (z=2.33, P=0.020) or 3-Hk treated males (z=3.71, P<0.001) possibly because distilled water, the vehicle for experimental injections, rehydrated both males injected with 3-Hk and sham males. Survival was not dependent on body size, since this variable was not retained by the best-supported model selected by Akaike Information Criterion methods (AIC).

**Figure 3.**
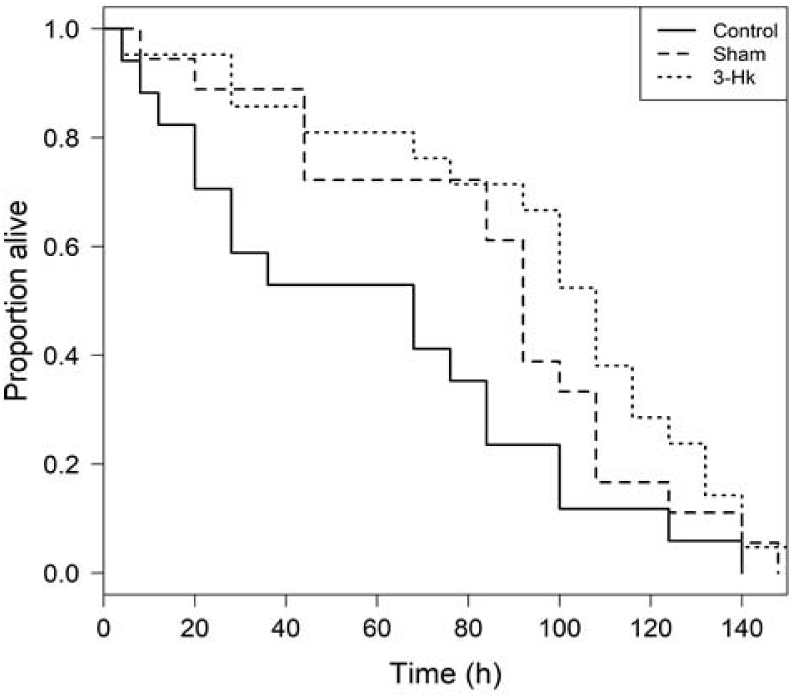
Survival time of *Hetaerina americana* males treated with 3-Hk, sham or control treatments.

In combinations with these results, we found that males treated with 3-Hk had higher values of *Rc* than males from the control or the sham treatments (Figure 4). Moreover, there was a significant effect (showed by the best AIC model) of treatments in interaction with survival time and in interaction with the RWS area (Sup. Table1). The interaction between treatment and survival shows that while red chroma was positively related to survival time in control males, red chroma was negatively related to survival time in sham males, and there was no relationship for 3-Hk treated males (see supplementary figure 2a). The interaction between treatment and RWS area indicated that even though RWS area was positively related to red chroma in all treatments, the slope was steeper in sham males than in control or 3-Hk treated males (see supplementary figure 2b).

**Figure 4.**
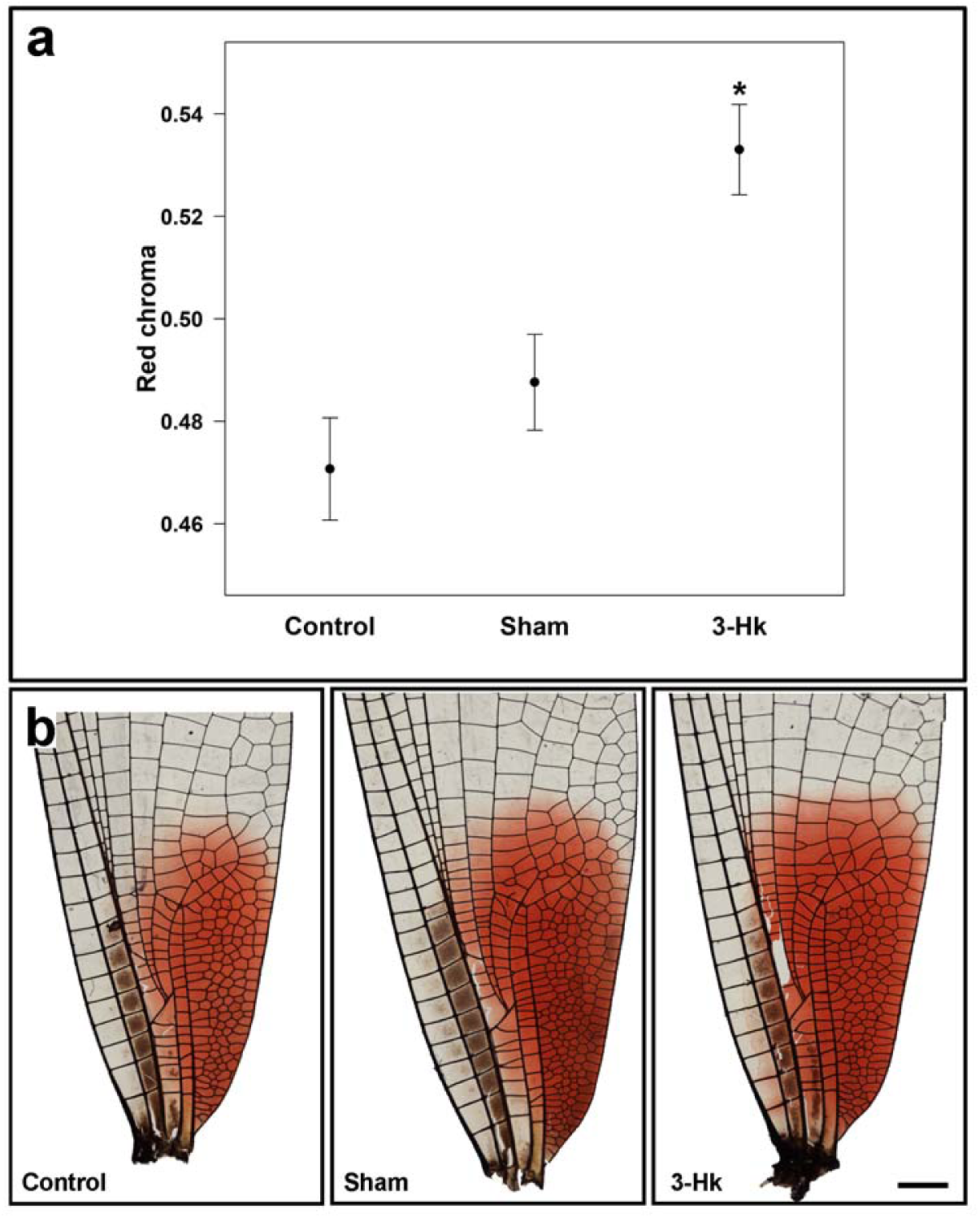
(a) Red chroma from RWS of *Hetaerina americana* males. Adult males injected with the toxic metabolite 3-Hk had higher values of red chroma than sham (i. e. Distiller water injected), and control (i. e. non-manipulated) males. (b) Bright field tile scan images of control, sham and 3-Hk treated males. Although the distribution of the wing veins is the same for all groups, qualitatively 3-Hk treatment shows stronger, diffuse red staining within the wing when compared with control and sham treatments. Scale bar: 1 mm. **\*** Significance at level of p < 0.01

On the other hand, mean values of *Rf* showed a slight difference between groups, with higher values in 3-Hk-treated males (1.09±0.17) than sham (0.79±0.18) and control males (0.84±0.20). Nevertheless, the only statistically significant selected predictor for *Rf* was RWS area, as males with larger red areas had higher values of *Rf* (F_1,28_=19.0, P<0.001; see supplementary figures 3a and 3b).

## Discussion

Ommochrome pigments were confirmed through all four of the biochemical properties we evaluated of the wings. Also, in support of this determination, TLC and spectral behaviour analyses indicated that xanthommatin and its reduced form, dihydroxanthommatin, are the autofluorescent ommochromes distributed along the wing spot veins (as shown by confocal microscopy; Fig. 2d and 2e). It is possible that ommatin D or rhodommatin are also present but that our pigment extraction method caused these pigments to spontaneously degrade into xanthommatin ^9,23^. TLC also revealed the presence of 3-Hk in the RWS. The participation of 3-Hk as a reddish pigment in insects is possible, as it was previously reported in red wing regions of lepidopterans ^23^ and in the cuticle of *Bombyx mori* ^24^. Nonetheless, its presence in *H. americana* wings was so dim that a slight band was hardly discernible (Fig.1b).

To our knowledge this is the first time that ommochromes have been observed by confocal microscopy techniques, taking advantage of the ommochromes’ property of autofluorescence when excited at a certain wavelength (561nm). This approach to reveal insect ommochromes could be complementary with other microscopy techniques previously used to observe these pigments, such as electronic microscopy, which has actually allowed the identification of the special organelles where ommochromes are produced, known as ommochromasomes ^18^.

In odonates, ommochromes have been found in the cuticle of some members of the dragonfly genera *Sympetrum* and *Crocothemis*, where these compounds are related to sexual dimorphism, maturation patterns and antioxidant capacity ^10^. Ommochromes in other taxa are also versatile pigments involved in several functions and not only colour production. Ommochromes are found in the eyes of insects ^18^ and to a lesser extent, in other structures such as the wings of some lepidopterans ^23^. Functionally, ommochromes are crucial in processing visual information in the insect compound eye ^9^, and they are suitable for transporting electrons or reacting with oxidants, reducers and free radicals ^25^. Notwithstanding, as evidenced by our survival experiment, tryptophan and its metabolites (including 3-Hk) are toxic at high concentrations in all insects ^9,26^. This dangerous situation can occurs due to important phenotypic changes during insects’ life cycles, for example, during molting ^9,26^. Ommochromes are also thought to be end-products of tryptophan detoxification ^9^. In fact, both 3-Hk and ommochrome pigments are found at high concentrations in the meconium of holometabolous insects, a waste product from the pupal stage ^19^. While some of these hemimetabolous insects, such as locusts, get rid of the toxic tryptophan by converting it to ommochromes and excreting it in faeces ^12^ storage of toxic byproducts in the form of ommochromes could be of particular relevance for detoxification in insects lacking pupation (i.e. hemimetabolous insects) such as our study animal and allied taxa. Nevertheless, this idea needs further testing.

Despite that our survival experiment shows that 3-Hk impairing survival when we estimated the LC_50_, this tryptophan metabolite affected the expression of the sexual trait in our animal model, resulting in an increment of red pigmentation in wings of male treated with 3-Hk compared to Control and Sham males. This result in combination with the lack of survival differences between Sham and 3-Hk males may support the idea that wing ommochromes may mitigate the toxicity of 3-HK. The increase in red pigmentation found in males treated with 3-Hk can be explained by several enzymatic and spontaneous reactions that may participate in a few metabolic processes, such as kynurenine-3-monooxygenase, which takes place during 3-Hk formation, or phenoxazone synthase, and has a role in ommochrome formation. Nonetheless, the physiological and molecular mechanisms that underlie the metabolic pathways from tryptophan metabolites to ommochromes are poorly known in biochemical terms to clarify ommochrome biosynthesis ^18^. Interestingly, our experimental manipulation with 3-Hk did not impact the area of the RWS. Previous works in our study species have provided evidence that this feature correlates with male energetic condition during aerial contests over mating territories ^21,27^. Thus, one explanation for our results is that by making the spot appear red, serves as the information signal that a territorial male need to convey to his conspecific and heteroespecific rivals during territorial tenure. This is compatible with a previous experiment in which the red spot was manipulated to appear blue (without manipulating spot area) in male territory holders ^28^. This change elicited high levels of aggression by rivals towards “blue spot” males, showing that it is red coloration but not spot area alone that is perceived as a the first signal of territoriality ^28^.

The honest signalling theory indicates only individuals in good condition (e.g. healthy) are able to afford the costs of generating and maintaining sexual signals ^29,30^. Food, parasites and free radicals are the main drivers of these signals ^30–33^. In this sense, our study indicates that ommochrome-pigmented sexual traits in *H. americana* males could act as honest indicator of nutritional condition, which is correlated with excretion ability. According to this, only males that acquire sufficient protein in their diets to require tryptophan detoxification and are able to use an effective detoxification mechanism to convert and excrete it as red ommochrome pigment will produce sufficient amounts of metabolic products to develop the red signal, which indicates this quality to rivals, and therefore gives them an advantage in acquiring and maintaining a mating territory. In this sense, the detoxification mechanism could have initially evolved because the production of ommochromes was an effective mechanism for dealing with metabolic waste products, and then became co-opted by sexual selection when these ommochromes were allocated to wings and indicated male nutritional condition to rivals. It is not that rivals assess the detoxification ability, but rather how much energy a rival has to utilize during aerial contests for territories. Given these metabolic processes and sexual selection mechanisms, individuals cannot “cheat” the system. This hypothesis—which we call here detoxifying ability signalling—should be tested in other pigments that are used in sexual selection contexts (one example is the case of butterfly pteridines ^34^). Interestingly, ommochromes in their reduced form are important antioxidants ^35^ and may combat oxidative stress, a situation that was recently shown in our study species ^36^. Thus, by reducing ommochrome toxicity, males may also benefit in terms of dealing with oxidative stress. This three-fold function – detoxification, antioxidant ability, and sexual signalling – provide support for a metabolic efficiency mechanism that could be informed through the expression of theses colourful pigments.

## Methods

### Animals

Adult male *H. americana* were captured in the riverine areas of the Tetlama River, Morelos, Mexico (18° 45’ 55’’N, 99° 14’ 45’’ W) with a butterfly net between 10.00 and 16.00h, the time at which males are most active ^37^. Males were aged according to three visual categories ^38^: 1) juvenile males, bearing soft, dorso-ventrally flexible, undamaged wings with a small and incompletely formed RWS; 2) sexually mature males, with less flexible wings, fully fixed red wing spot and no signs of thoracic pruinescence (Fig. 1a), and 3) old males, with abundant thoracic pruinescence and inflexible, damaged or worn wings with the same RWS size as sexually mature males. Given that animals from the second category have a better physiological condition and will remain alive for a longer period from the time of capture ^28^, we only used sexually mature males.

#### 1. Presence of ommochromes in the RWS of *H. americana* males

##### 1.1. Pigment extraction

To determine whether ommochrome pigments are present in the RWS, we sacrificed 30 adult males by freezing for 20 minutes at −20°C soon after collection. These animals were then dried in a drying oven (Heratherm-ThermoFisher) for 24 hours at 30°C. The four wings were removed from each dried male, but only the RWS from the forewings was used. To extract the red pigment from the forewings we used the methodology proposed by Nijhout^23^, with some modifications from Riou and Christidès ^11^. All spots were pooled and homogenized in methanol at 4°C and centrifuged for 5 minutes at 14,000 g. Then, the precipitate was washed twice with 99% methanol and three times with ethyl ether by repeated suspension and centrifugation (5 minutes at 14,000g). After the ethyl ether was evaporated from the final precipitate, 4 ml of acidic methanol (100% methanol plus 0.5% hydrochloric acid) were added to the residue and the suspension was centrifuged for 10 minutes at 14,000 g. After centrifugation, the supernatant was concentrated to one-quarter of its volume using a vacuum concentrator (SpeedVac-Thermo®, Mod. ISS110). We added 2 ml of distilled water and SO gas to this solution and, after incubating overnight for 8 hours at 2 °C, the red pigment obtained was precipitated by centrifugation for 10 min at 14,000 g and washed with a 2 mL of cold water. The sample was then dried in a drying oven (Heratherm-ThermoFisher) at 28°C until water evaporated. The powder obtained was used to determine the nature of the red wing pigment.

##### 1.2. Pigment determination

We performed four complementary methods to evaluate presence of ommochromes via their biochemical properties: Redox behaviour, spectral absorbance, thin layer chromatography and autofluorescence by confocal microscopy.

###### 1.2.1. Redox behaviour and spectral absorbance

The most conspicuous property of ommochromes is their reduction-oxidation behaviour (i.e. Redox), which is observed by a colour change from red when reduced to yellow when oxidized ^9^. Hence, we evaluated whether our red powder changed colour when it was subjected to reductant or oxidant conditions. The red spot powder was dissolved in phosphate buffer (0.067 M, pH 8.5) containing 10 μL of 1% ascorbic acid as a reductant component to observe the resulting colour. Next, we added 10 μL of 1% NaNO_2_ as an oxidant component and the colour changes were observed. Finally, we again added 10 μL of 1% ascorbic acid to test whether the colour reverted to its previous state.

Complementary with this Redox behaviour, all ommochromes have a particular spectral behaviour, which is characterized by 2 to 4 spectral peaks, distributed throughout the UV and visible ranges of the spectrum ^9,39^. These spectral peaks have been traditionally characterized when ommochromes are dissolved in both phosphate buffer (pH 7.0) and 5N HCl ^23,40^. Therefore, to determine whether our red powder showed the typical spectral behaviour of ommochromes, we measured the spectral peaks when the red powder was dissolved in these solvents. To accomplish this, we prepared two different solutions using 20 µg of the red powder and 100 µL of phosphate buffer or 5N HCl. Given that spectral peaks of the ommochrome xanthommatin are the best characterized by previous studies ^9,18^, we processed the samples to convert any ommochromes that could be present in the RWS powder (e.g. ommatin D or dihydroxanthommatin) into xanthommatin. This conversion allows us to ensure the specificity in the resulting peaks. For this purpose, in the case of the phosphate buffer, the solution was refrigerated at 4°C for 12 hours, while the 5N HCl solution was kept at 25 °C in a drying oven (Heratherm-ThermoFisher) for 5 days ^40^. After this period, both solutions were centrifuged (14,000 g for 5 min) and the absorbance of the supernatant was measured from 220-600 nm wavelength using a spectrophotometer (Hitachi, Mod. U-3900). Finally, we compared the spectral peaks obtained in both solvents with those described by Nijhout ^23^, who used the same solvents to evaluate spectral peaks of ommochromes extracted from the ventral hind wing pattern of the *linea* and *rosa* forms of the lepidopteran *Precis coenia* ^23^.

###### 1.2.2 Thin layer chromatography (TLC)

TLC is a technique that allows the separation and characterization of ommochromes from insects when these pigments are present ^23^. For this purpose, we used a 10×20 cm silica gel plate (Merck, Darmstadt, Germany, Silica Gel 60 F254). This plate was divided into 4 channels (2 cm between channels). The division was made from the narrower side of the plate (10 cm) and a thin base line was drawn with a pencil at 1 cm from the bottom of the plate in each channel. At this base line, we added 10 μL from the wing spot red powder that was previously diluted in acidic methanol (1 mL ^23^), and the plate was left in a vertical position in a chromatographic chamber (Aldrich). The chamber was previously saturated with a developing solvent of phenol:water (3:1 by volume) using a small piece of filter paper (JM 3639, JoyLab) around the chamber. The silica gel plate was left inside the chamber for 2 hours to allow the complete separation of pigments. After this time, we identified the bands formed along the plate in both ultraviolet and visible light spectrum using an UV/White transilluminator (Safe Imager 2.0, Invitrogen). Then, we measured the distance (mm) at which these bands were located with respect to the base line as well as the distance travelled by the developing solvent in each channel, and we obtained an average value for each band and for the solvent. These distances were used to obtain the retention factor (R_f_) for each pigment found. Finally, to determine whether any R_f_ obtained in our sample corresponded to ommochrome pigments, these values were compared with the R_f_ of authentic synthetic ommochromes previously reported from the same silica gel/phenol TLC system ^23^, since there are no commercially available ommochrome standards available to date ^18^.

###### 1.2.2. Autofluorescence analysis

The presence of indole groups in the chemical structure of all ommochromes gives them the property of autofluorescence at certain wavelengths ^9^. This biochemical feature is a helpful way to determine the nature of certain colours in insects, since only red pigments derived from ommochromes and aphins (i.e. a pigment only present in aphids and not in any other insects ^41^) will show this property ^9,42^. Therefore, one last complementary test to determine whether ommochromes are present in the RWS of *H. americana* was to observe autofluorescence induced by excitation with specific laser lines and evaluated with a spectral detector using multiphoton microscopy. To accomplish this, five forewings from adult males were carefully mounted onto pre-cleaned microscope slides (Lauka, CDMX, México) and covered with 0.17mm thick 180 x 180 mm coverslips. Then, the edges of the coverslip were carefully sealed to the slide with nail hardener and slides were kept at room temperature until imaging. Bright field tile imaging was carried out with an upright microscope (Olympus BX51-WI, Olympus Corporation, Tokyo, Japan) equipped with an XYZ motorized stage (MAC6000, Ludl Electronic Products, Ltd., Hawthorne, NY, USA), an RGB CCD camera (MBF CX9000, MBF Bioscience, Williston, VT, USA) and StereoInvestigator software (v. 9.0.1, MBF Bioscience). Imaging was done using a UPLAN FL N 10X N.A. objective. To visualize the autofluorescence produced by the red wing spot, we acquired XYZλ images from both pigmented and non-pigmented regions of the same wing with a Nikon A1R^+^ laser confocal scanning head coupled with an Eclipse Ti-E inverted microscope (Nikon Corporation, Tokyo, Japan) equipped with a motorized stage (TI-S-E, Nikon). We thus excited the samples using four lasers of different wavelengths: 405 (2mW), 488 (70mW), 561 (1.4mW) and 647 nm (1.25mW). We evaluated the resulting signals using a 32-channel spectral detector (10 nm resolution, from 425 to 735 nm wavelength) plus a transmitted light detector. The pinhole value was set at 29.4 µm for all lasers. Images were captured with NIS Elements C software v. 4.50 (Nikon); and XYZ□ resulting images were converted to single-plane images by applying maximum intensity projections (MIP) in order to show the brightest fluorescence information for all Z planes for all wavelengths at a glance with the same software.

Finally, fluorescence single plane tile imaging of the forewings was done with a CFI Plan Fluor 10X N.A. 0.3 objective, using 7mW of 561nm laser power, pinhole aperture of 195.4µm, and a GaAsP detector, all controlled through NIS Elements C software v4.50 (Nikon).

#### 2. Toxic effects of 3-Hk in *H. americana* males

Given that 3-Hk has been reported to be fatal to insects ^15^, we evaluated whether this substance kills *H. americana* males. We performed an experiment in captivity to calculate the median lethal concentration (LC_50_) of 3-Hk. For this, 3-Hk (Sigma-Andrich; Catalogue number: 2147-61-7) was dissolved in distilled water using a vortex mixer for 10 minutes. Five different 3-Hk concentrations were injected into 75 adult males (15 per group): 0 (only distilled water), 1, 100, 1000 and 10,000 μg mL^−1^. The injection took place in the dorsal thoracic region, where the wings are inserted. Animals received 4µl of each 3-Hk concentration using a microsyringe (10 µL, Hamilton 80330; Hamilton, Reno, Nevada). Individual manipulation lasted no more than 1 min. Males were then placed individually into transparent plastic 5-mL assay tubes (Simport, Canada) with a piece of wood as a perch, moist pieces of cotton to provide humidity and a temperature of 26 ◦C inside the tubes. Males were not fed during the experiment and conditions of captivity were always the same. Twenty-four hours after injection, male mortality was recorded for each of the five groups. This survival experiment concluded that 3-Hk is toxic for adult males of *H. Americana*; the LC_50_ we determined for this species which was estimated at 368.69μg mL^−1^ (C.I 95%: 78.5-1729.9μg mL^−1^; See results section for further details).

#### 3. Mitigation of 3-Hk toxic effects by deposition of ommochromes in RWS

To evaluate whether males are able to counteract the toxic effect of 3-Hk by depositing ommochromes into their wings, forming the red wing spots, we performed another captivity experiment administrating 3-Hk in males, then determining whether the ommochromes were subsequently deposited into their RWS. To accomplish this, 54 adult males were captured and randomly allocated to three different treatments: 1) 3-Hk treatment (N=20 males injected with 363µg of 3-Hk diluted in 4 µL of PBS 1x—the previously estimated LC_50_), 2) Sham treatment (N=17 males injected with 4 µL of PBS 1X), and 3) Control treatment (N=17 males with no manipulation). After manipulation, males were monitored in captivity every 4 h to record the time to death, and the experiment ended when the last male died. Captivity conditions were the same as those used to calculate the LC_50_. After the last male died, the anterior wings from all individuals were removed from the body. The effect of 3-Hk on wing red pigmentation was evaluated by two complementary techniques: 1) quantification of red chroma (*Rc*) by spectrophotometry (see also ^21^), and 2) the relative fluorescence *(Rf)* by confocal microscopy. The red chroma of RWS from the different treatments was calculated as the proportion of total reflectance (from 360 to 740 nm) occurring in red wavelengths (600-700 nm) using a spectrophotometer (MINOLTA CR-200, Konica Sensing Inc., Osaka, Japan; for a similar procedure see ^28^).

*Rf* was determined from confocal XYZ images obtained from a random sample of 10 forewings from males of each treatment. Laser scanning confocal microscope Z-stack images (512×512 pixels, 12-bits, 3 µm interval) were acquired with a CFI Plan Fluor, 10X, N.A. 0.3 objective, using 1.4mW of 561nm laser power for excitation, pinhole aperture of 20.43 µm, emission filter 595/50, and a GaAsP detector, all controlled through NIS Elements C software v4.50 (Nikon). Z-images were then processed using Image J software ^43^, performing a Z-projection with the pixels obtained from maximum intensity projection (MIP) as the reference. The MIP image was converted from 12 to 8-bits and a histogram of the pixel intensity value was extracted. *Rf* value from each Z-projection was calculated by multiplying the intensity value of the histogram with its corresponding number of pixels then dividing by the total number of pixels present in each image (262144 pixels ^43^). Finally, *Rf* value of all Z-projections were averaged to obtained a unique *Rf* value per individual.

#### 4. Statistical analyses

To determine whether ommochromes are responsible for the male RWS we qualitatively analyzed the biochemical properties previously mentioned: redox behaviour, spectral absorbance, *Rf* of purified ommochromes, and autoflourescence. To evaluate whether 3-Hk, the precursor of ommochrome pigments, is a toxic tryptophan metabolite for adult males, we performed a survival analysis to calculate the LC_50_ for these males through the Trimmed Spearman-Karber method ^44^. We included in the model the following predictors: treatment (3-Hk, Sham, and Control), wing length, RWS area, and their interactions. Analysis was done in R ^45^ using the tsk package ^46^.

To evaluate whether males of *H. americana* are able to counteract the toxic effect of 3-Hk by depositing ommochrome in their RWS, we evaluated both the survival differences between treatments (i.e. 3-Hk, sham, and control) and the differences in colour properties of *Rc* and *Rf*. Survival after treatment was evaluated with a Cox proportional hazard regression, whose predictors were the additive effects of treatment and wing length (a proxy of body size ^28^. This model was simplified based on AIC values to obtain the best supported model (i.e. the model with the lowest AIC-value).

To determine differences in *Rc* and *Rf* we used linear mixed and linear models respectively. For *Rc* the predictor variables were treatment, survival time, wing length, RWS area and the interactions treatment*survival time, treatment*wing length and treatment*RWS. Given that *Rc* was measured for both the left and right forewings of each damselfly, individual identity was included as a random effect in this analysis. For the case of *Rf* we used the same predictors as for *Rc.* The initial models were reduced based on AIC and the best supported model is reported. *P*-values of the predictor variables and interactions were obtained using likelihood ratio tests for *Rc* and with F tests for *Rf*. Variance homogeneity was tested using the Fligner-Killeen test, normality of residuals was inspected visually from normal q-q plots and the presence of outliers was evaluated with Cook’s distances (none were found—all Cook’s distances<1). All analyses were done in R software ^45^ according to Crawley ^47^ and Zuur and collaborators ^48^.

## Data availability

All data are available from the corresponding author on reasonable request.

## Acknowledgements

We are grateful to Ana Rivas-Caicedo for her advice on the microscopy techniques implemented in the present study. We thank Roberto Arreguín Espinosa to his advice on pigment determination. We thank Fernanda Baena for helpful discussion and fieldwork. We also thank Lynna Marie Kiere for her style revision and thoughtful comments, and Osiris Gaona for helpful discussion. This project was funded by UNAM-PAPIIT [grant numbers IA207019 and IN204610] and CONACYT Ciencia Básica [grant number 241744].

## Author contributions

I.G.-S conceived this study. All authors designed this study. I.G.-S., and D. G.-T. collected fieldwork data, M.T.-R. collected and analyzed microscopy data, I.G.-S. and D. G.-T. analyzed all data. All authors contributed in writing the first draft and approved the final version of the manuscript.

## Additional information

Supplementary information is available for this paper upon correspondence and requests for materials should be addressed to I.G.-S or A.C.-A.

---

**Figure S1a.**
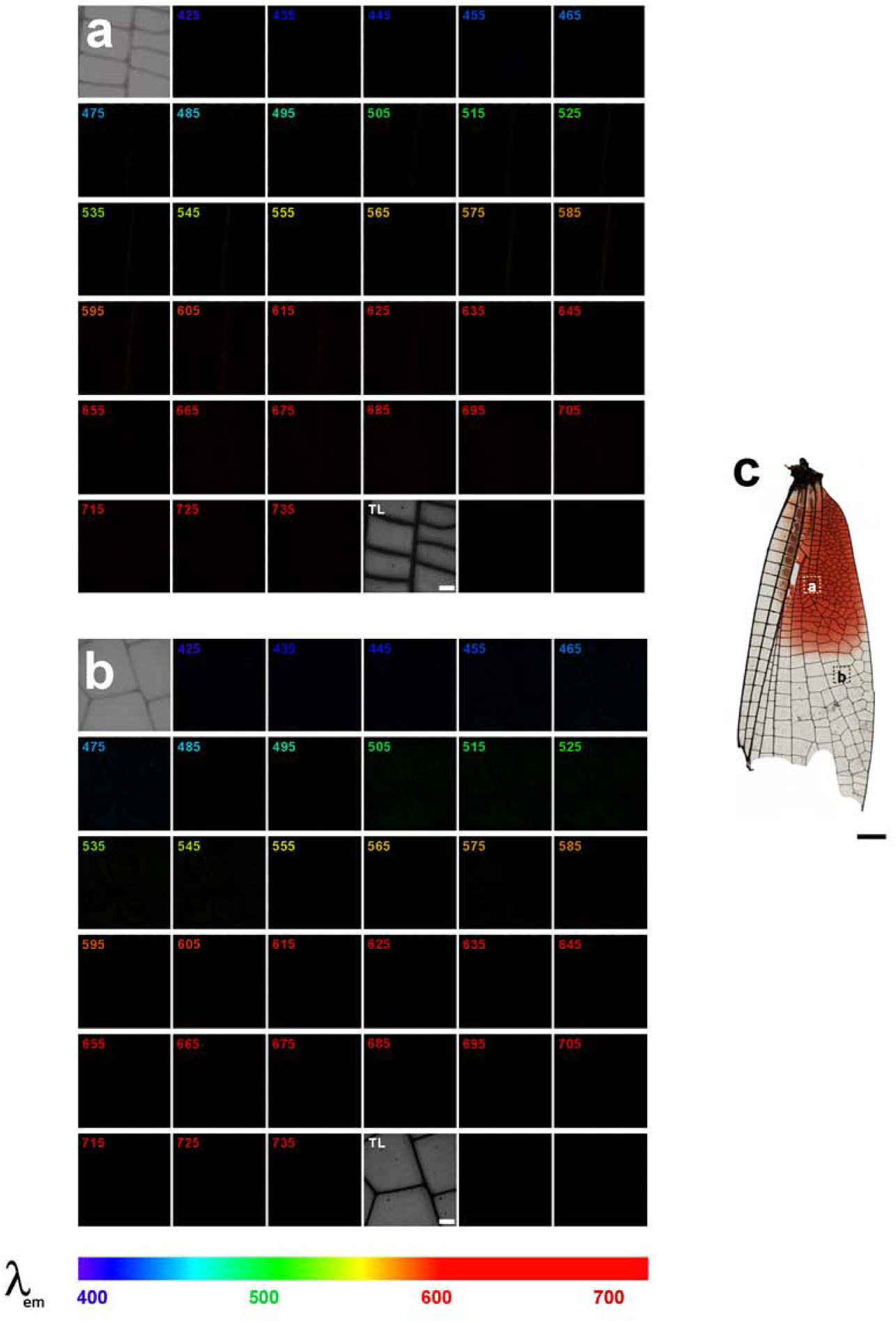
Fluorescence XYZ□ scans (425-735 nm) of *Hetaerina americana* wing excited at 405 nm. Maximum intensity projection images of (a) a sub-region of the red-pigmented area, and (b) a sub-region of the transparent area, depicted in (c). A relatively weak fluorescence intensity was found in wing veins of the anterior region from 505 to 725 nm (a). In contrast, no apparent signal is present in wing veins but in the wing between 465-615 nm in the medial region of dragonfly wing (b). Scale bar: 100 □ m.

**Figure S1b.**
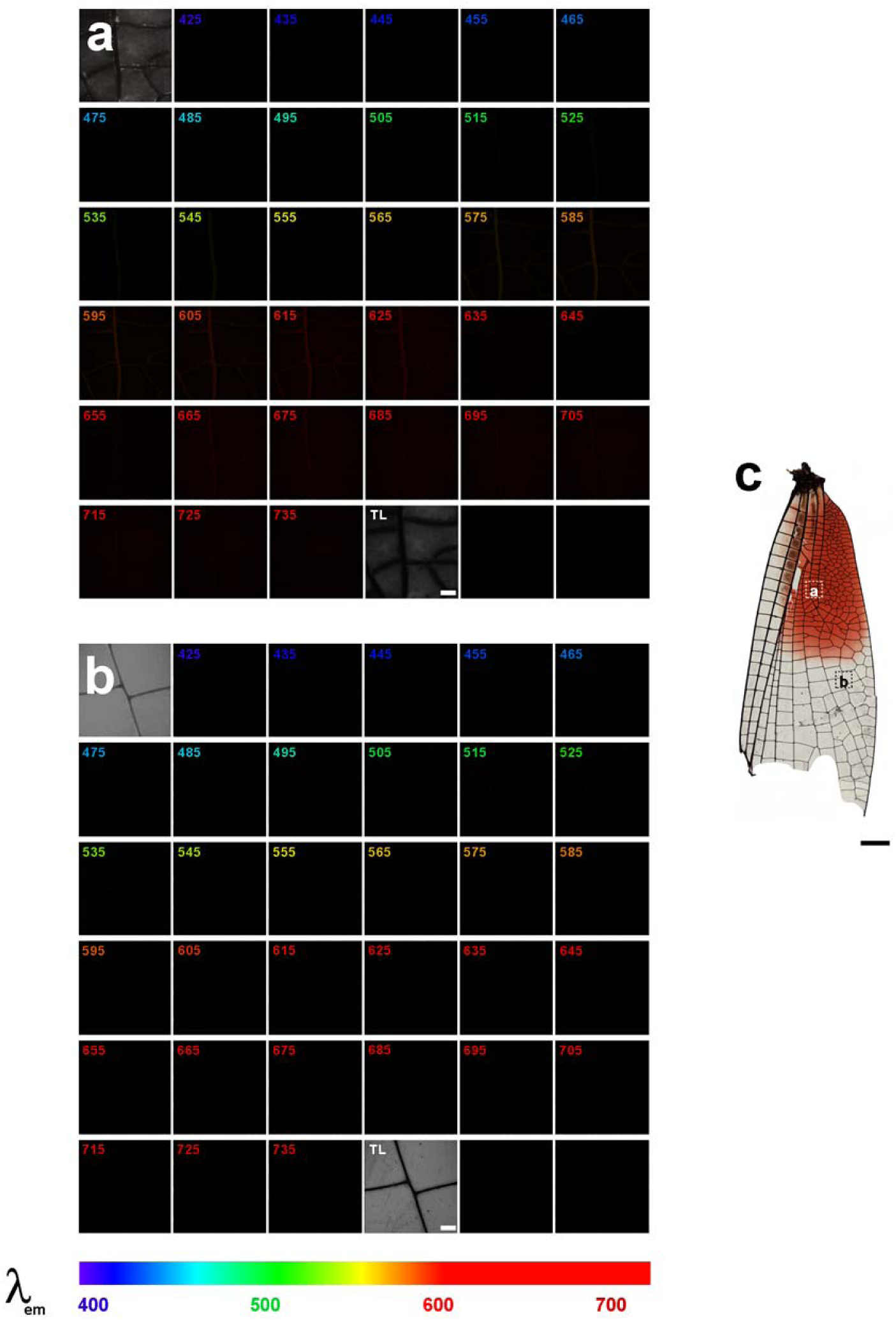
Fluorescence Zλ scans (425-735 nm) of *Hetaerina americana* wing excited at 488 nm. Maximum intensity projection images of (a) a sub-region of the red-pigmented area, and (b) a sub-region of the transparent area, depicted in (c). A relatively weak fluorescence intensity was found in the wing veins of the anterior region from 505 to 735 nm (a). In contrast, no apparent signal is present in the medial region of the damselfly wing (b). Scale bar: 100 μm.

**Figure S1c.**
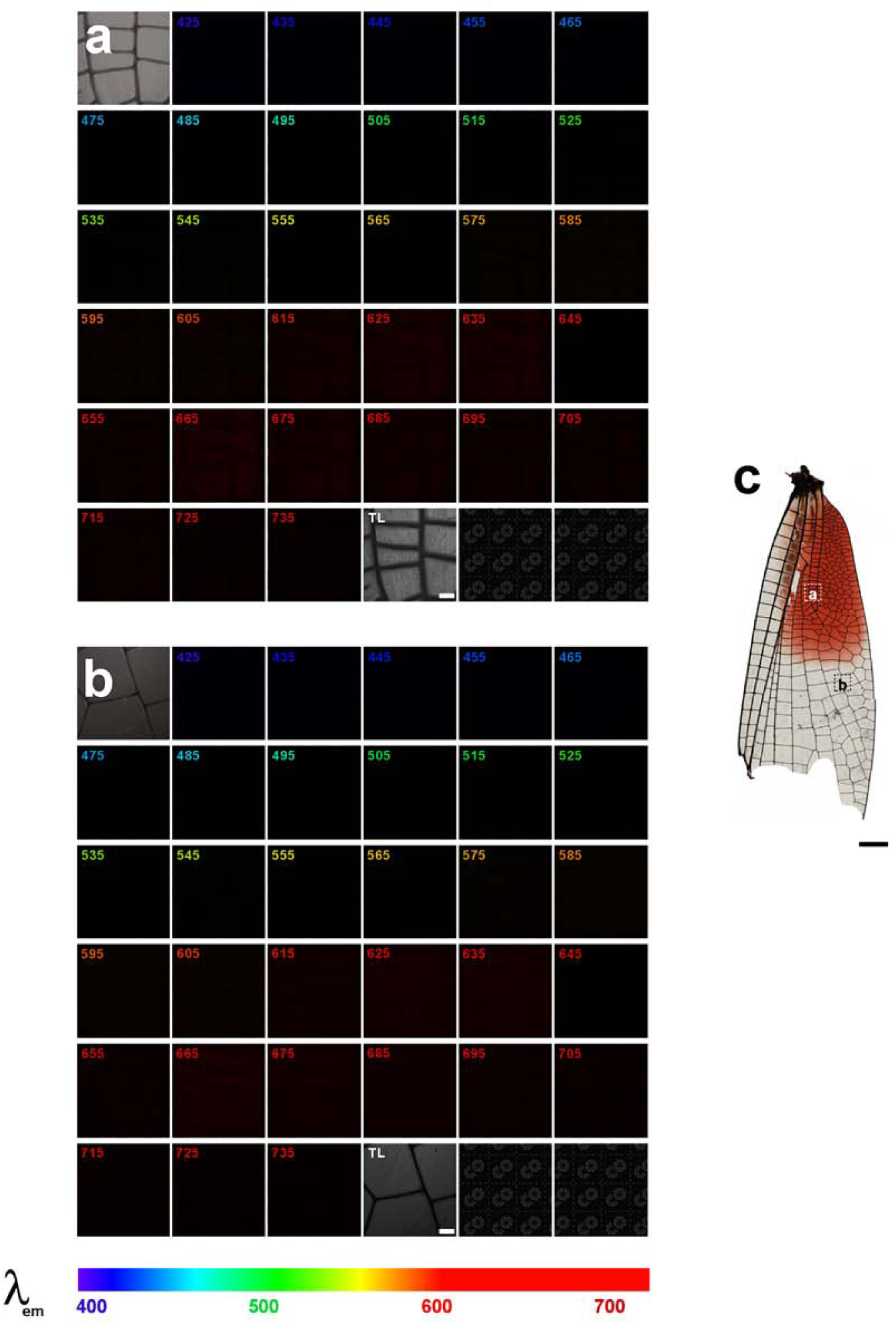
Fluorescence Zλ scans (425-735 nm) of *Hetaerina americana* wing excited at 646 nm. Maximum intensity projection images of (a) a sub-region of the red-pigmented area, and (b) a sub-region of the transparent area, depicted in (c). There was no apparent signal in wing veins but curiously a relatively weak fluorescence intensity was found in the wing of both anterior (a) and medial (b) regions from 575 to 735 nm. Scale bar: 100 μm.

**Figure S2.**
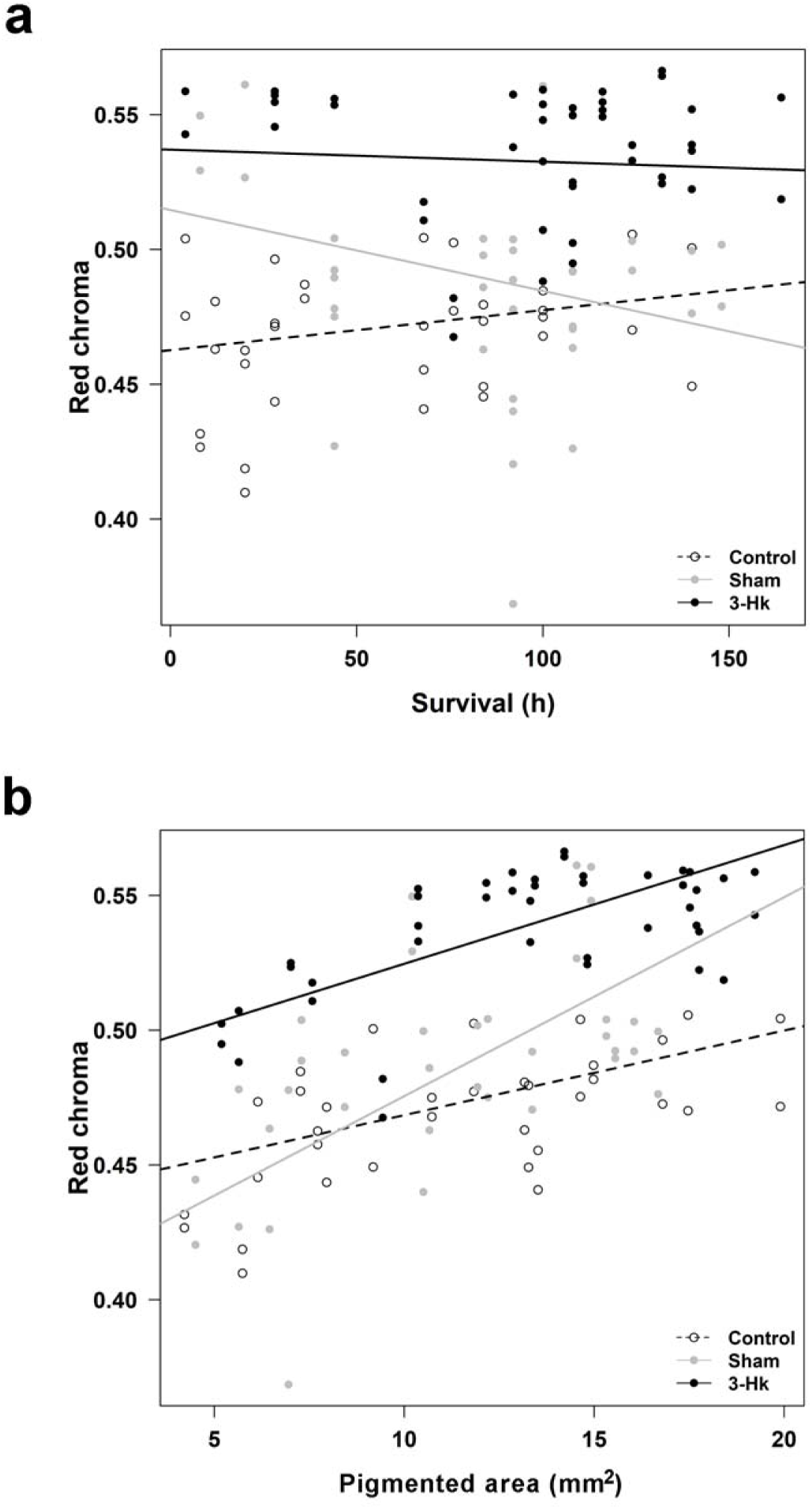
(a) Relationship between survival time and red chroma in each experimental treatment. While in males treated with 3-Hk(continuous black line) red chroma was not affected by survival time, control males (black dotted line) showed a positive relationship. In sham males (gray line), chroma was also affected by survival but in a negative direction. (b) Red chroma was also affected by RWS area. All treatments showed a positive relationship, but with different slopes.

**Figure S3.**
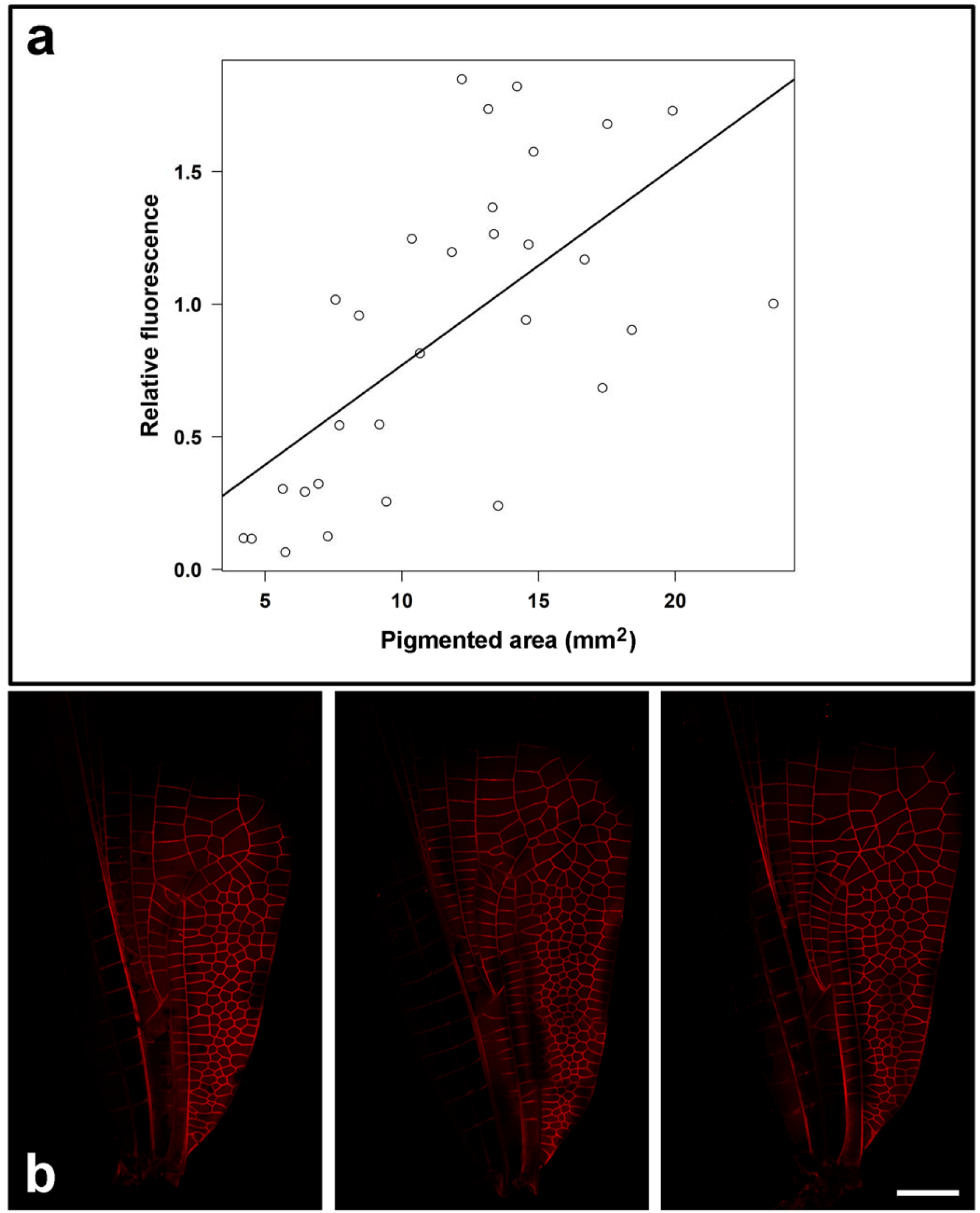
(a) Fluorescence XYZλ scans (425-735 nm) of *H. americana* wings excited at 561 nm of control, sham and 3-Hk males. Although 3-Hk males showed higher fluorescence values, the only significant predictor for this colour property was RWS area, which had a positive relationship in all treatments (b).

**TABLE S1.** Linear mixed effect model to explain variation in wing red chroma in *Hetaerina americana* males manipulated with 3-Hk, sham or control treatments. Significant predictors are shown in bold. L. Ratio=Likelihood ratio. NS=Not selected in the best supported model.

| Effect on red chroma | L. Ratio | P-value |
| --- | --- | --- |
| Treatment | 69.07 | **<0.001** |
| Survival time | 8.97 | **0.030** |
| Wing length | 9.99 | **0.019** |
| Pigmented area | 43.42 | **<0.001** |
| Treatment X Survival | 8.29 | **0.016** |
| Treatment X Wing length | NS | NS |
| Treatment X Pigmented area | 8.43 | **0.014** |

## References

1. Cochran, D. Insect biochemistry and function. (Springer-Science business media, 1975).

2. Timmerman, S. & Berenbaum, M. Uric acid deposition in larval integument of black swallowtails and speculation on its possible functions. J. Lepid. Soc. 53, 104–107 (1999).

3. Andersson, M. Sexual selection. (1994).

4. Zera, A. J. & Harshman, L. G. The physiology of life history trade-offs in animals. Annu. Rev. Ecol. Syst. 32, 95–126 (2001).

5. Vukusic, P., Chittka, L. & Chapman, R. F. Visual signals: color and light production. in The Insects:structure and function (eds. Chapman, R., Simpson, S. & Douglas, A.) 793–823 (Cambridge University Press, 2013). doi:10.1017/cbo9781139035460.032.

6. von Schantz, T., Bensch, S., Grahn, M., Hasselquist, D. & Wittzell, H. Good genes, oxidative stress and condition-dependent sexual signals. Proceedings. Biol. Sci. 266, 1–12 (1999).

7. McGraw, K. J. The antioxidant function of many animal pigments: are there consistent health benefits of sexually selected colourants? Anim. Behav. 69, 757–764 (2005).

8. Monaghan, P., Metcalfe, N. B. & Torres, R. Oxidative stress as a mediator of life history trade-offs: mechanisms, measurements and interpretation. Ecol. Lett. 12, 75–92 (2009).

9. Linzen, B. The Tryptophan → Ommochrome Pathway in Insects. Advances in Insect Physiology 117–246 (1974) doi:10.1016/s0065-2806(08)60130-7.

10. Futahashi, R., Kurita, R., Mano, H. & Fukatsu, T. Redox alters yellow dragonflies into red. Proc. Natl. Acad. Sci. U. S. A. 109, 12626–12631 (2012).

11. Riou, M. & Christidès, J.-P. Cryptic Color Change in a Crab Spider (*Misumena vatia*): Identification and Quantification of Precursors and Ommochrome Pigments by HPLC. J. Chem. Ecol. 36, 412–423 (2010).

12. Chapman, R. The Insects: Structure and Function. (Cambridge University Press, 1998).

13. Kayser, H. Ommochrome formation and kynurenine excretion in *Pieris brassicae*: Relation to tryptophan supply on an artificial diet. J. Insect Physiol. 25, 641–646 (1979).

14. Connolly, K., Burnet, B. & Sewell, D. Selective Mating and Eye Pigmentation: An Analysis of the Visual Component in the Courtship Behavior of *Drosophila melanogaster*. Evolution (N. Y). 23, 548 (1969).

15. Cerstiaens, A. et al. Neurotoxic and neurobehavioral effects of kynurenines in adult insects. Biochem. Biophys. Res. Commun. 312, 1171–1177 (2003).

16. Oxenkrug, G. F. The extended life span of *Drosophila melanogaster* eye-color (white and vermilion) mutants with impaired formation of kynurenine. J. Neural Transm. 117, 23–26 (2010).

17. Oxenkrug, G. F., Navrotskaya, V., Voroboyva, L. & Summergrad, P. Extension of life span of *Drosophila melanogaster* by the inhibitors of tryptophan-kynurenine metabolism. Fly (Austin). 5, 307–309 (2011).

18. Figon, F. & Casas, J. Ommochromes in invertebrates: biochemistry and cell biology. Biol. Rev. 94, 156–183 (2019).

19. Ogawa, H. & Hasegawa, K. Kynureninase and its activity during metamorphosis of the silkworm, *Bombyx mori*. Insect Biochem. 10, 589–593 (1980).

20. Hooper, R. E., Tsubaki, Y. & Siva-Jothy, M. T. Expression of a costly, plastic secondary sexual trait is correlated with age and condition in a damselfly with two male morphs. Physiol. Entomol. 24, 364–369 (1999).

21. Contreras-Garduno, J., Buzatto, B. A., Serrano-Meneses, M. A., Najera-Cordero, K. & Cordoba-Aguilar, A. The size of the red wing spot of the American rubyspot as a heightened condition-dependent ornament. Behav. Ecol. 19, 724–732 (2008).

22. Córdoba-Aguilar, A. & González-Tokman, D. M. The Behavioral and Physiological Ecology of Adult Rubyspot Damselflies (*Hetaerina*, Calopterygidae, Odonata). Advances in the Study of Behavior 311–341 (2014) doi:10.1016/b978-0-12-800286-5.00007-9.

23. Nijhout, H. Ommochrome pigmentation of the linea and rosa seasonal forms of *Precis coenia* (Lepidoptera: Nymphalidae). Arch. Insect Biochem. Physiol. 36, 215–222 (1997).

24. Meng, Y., Katsuma, S., Mita, K. & Shimada, T. Abnormal red body coloration of the silkworm, *Bombyx mori*, is caused by a mutation in a novel kynureninase. Genes to Cells 14, 129–140 (2009).

25. Needham, A. The Significance of Zoochromes. (Springer, 1974).

26. Manoukas, A. Effect of excess levels of individual amino acids upon survival, growth and pupal yield of *Dacus oleae* (Gmel.) larvae. Zeitschrift für Angew. Entomol. 91, 309–315 (1981).

27. Grether, G. F. Survival cost of an intrasexually selected ornament in a damselfly. Proc. R. Soc. London. Ser. B Biol. Sci. 264, 207–210 (1997).

28. González-Santoyo, I., González-Tokman, D. M., Munguía-Steyer, R. E. & Córdoba-Aguilar, A. A mismatch between the perceived fighting signal and fighting ability reveals survival and physiological costs for bearers. PLoS One 9, e84571–e84571 (2014).

29. Zahavi, A. Mate selection—A selection for a handicap. J. Theor. Biol. 53, 205–214 (1975).

30. Hamilton, W. & Zuk, M. Heritable true fitness and bright birds: a role for parasites? Science (80-.). 218, 384–387 (1982).

31. Sheldon, B. C. & Verhulst, S. Ecological immunology: costly parasite defences and trade-offs in evolutionary ecology. Trends Ecol. Evol. 11, 317–321 (1996).

32. Montoya, B., Valverde, M., Rojas, E. & Torres, R. Oxidative stress during courtship affects male and female reproductive effort differentially in a wild bird with biparental care. J. Exp. Biol. 219, 3915–3926 (2016).

33. Olzer, R., Ehrlich, R., Heinen-Kay, J., Tanner, J. & Zuk, M. Reproductive behavior. in Insect Behavior: from mechanisms to ecological and evolutionary consequences 189–202 (2018).

34. Tigreros, N., Mowery, M. A. & Lewis, S. M. Male mate choice favors more colorful females in the gift-giving cabbage butterfly. Behav. Ecol. Sociobiol. 68, 1539–1547 (2014).

35. Futahashi, R. Color vision and color formation in dragonflies. Curr. Opin. Insect Sci. 17, 32–39 (2016).

36. Martínez-Lendech, N., Golab, M. J., Osorio-Beristain, M. & Contreras-Garduño, J. Sexual signals reveal males’ oxidative stress defences: testing this hypothesis in an invertebrate. Funct. Ecol. 32, 937–947 (2018).

37. Contreras-Garduño, J., Canales-Lazcano, J. & Córdoba-Aguilar, A. Wing pigmentation, immune ability, fat reserves and territorial status in males of the rubyspot damselfly, *Hetaerina americana*. J. Ethol. 24, 165–173 (2006).

38. Plaistow, S. & Siva-Jothy, M. Energetic constraints and male mate-securing tactics in the damselfly *Calopteryx splendens* xanthostoma (Charpentier). Proc. R. Soc. London. Ser. B Biol. Sci. 263, 1233–1239 (1996).

39. Bolognese, A., Liberatore, R., Riente, G. & Scherillo, G. Oxidation of 3-hydroxykynurenine. A reexamination. J. Heterocycl. Chem. 25, 1247–1250 (1988).

40. Umebachi, Y. & Uchida, T. Ommochromes of the testis and eye of *Papilio xuthus*. J. Insect Physiol. 16, 1797–1812 (1970).

41. Shamim, G., Ranjan, S. K., Pandey, D. M. & Ramani, R. Biochemistry and biosynthesis of insect pigments. Eur. J. Entomol. 111, 149–164 (2014).

42. Insausti, T. C. & Casas, J. The functional morphology of color changing in a spider: development of ommochrome pigment granules. J. Exp. Biol. 211, 780–789 (2008).

43. Abramoff, M., Magalhaes, P. & Ram, S. Image processing with ImageJ. Biophotonics Int. 11, 36–42 (2004).

44. Hamilton, M. A., Russo, R. C. & Thurston, R. V. Trimmed Spearman-Karber method for estimating median lethal concentrations in toxicity bioassays. Environ. Sci. Technol. 11, 714–719 (1977).

45. R Core Team. R: A Language and Environment for Statistical Computing. (2019).

46. Stone, B. R. tsk: Trimmed Spearman-Karber Method. (2012).

47. Crawley, M. J. The R Book Second Edition. (Wiley, 2012).

48. Zuur, A. F., Ieno, E. N., Walker, N., Saveliev, A. A. & Smith, G. M. Mixed effects models and extensions in ecology with R. Statistics for Biology and Health (2009) doi:10.1007/978-0-387-87458-6.

